# Categorical Color Structure Persists in Color Vision Deficiency

**DOI:** 10.64898/2026.09.13.751220

**Authors:** Aimee Martin, Doris I. Braun, Karl R. Gegenfurtner

## Abstract

Color vision deficiencies affect color perception and discrimination, but their consequences for the structure and stability of color categories remain less clear. We examined how trichromats, anomalous trichromats, and dichromats assigned 450 physical Munsell color samples to eleven basic color categories and measured the stability of these assignments in a retest two hours later. In each session, participants sorted a randomly mixed set of chips by hand and selected one representative chip for each category. We quantified agreement within and between color-vision groups as well as test-retest reliability within participants.

For the 217 chips that all trichromats assigned to the same category, anomalous trichromats used the same category label for 85% of chips, and their category assignments were less stable across sessions as those of trichromats. Within anomalous trichromats poorer red-green discrimination, indexed by higher RG-CAD scores, was associated with lower sorting consistency. Dichromats showed lower agreement with trichromats and anomalous trichromats and greater prototype instability, but their color categories remained systematically organized rather than random. Information-theoretic analyses revealed a graded increase in categorical uncertainty from trichromats through anomalous trichromats to dichromats together with increasing reliance on lightness.

Thus, the principal effect of color vision deficiency was not a collapse of color categories, but a graded reduction in the precision with which different shades were assigned to otherwise recognizable categories. In a separate subset of nine anomalous trichromats, EnChroma glasses produced no measurable immediate shift toward trichromatic category structures. Overall, our findings suggest that categorical color representations in anomalous trichromats and dichromats remain remarkably robust despite substantial sensory deficits.

## Introduction

Inherited red-green color vision deficiency (CVD) affects the sensitivity to colors of a notable proportion (6%-8%) of the male population ([1]; for reviews, see [2–4]). Depending on the alteration or loss of photopigment genes two broad kinds of deficiency are identified: anomalous trichromacy, and dichromacy ([5,6]; for reviews, see [7,8]). In anomalous trichromacy, all three photoreceptor cone types are functional, but the spectral sensitivity of one cone type is shifted [9]. In deuteranomalous observers, the most prevalent form of red–green CVD (approximately 6% of males), the sensitivity curve of the medium-wavelength (M) cones is shifted toward that of the long-wavelength (L) cones, whereas in protanomalous observers the sensitivity of the L cones is shifted toward that of the M cones. In dichromacy, one cone class is entirely absent or not functional: deuteranopes lack M cone and protanopes lack L cone inputs, leading to more profound confusions along the red–green axis (for review, see [2]).

Research on CVD has historically emphasized sensory discrimination deficits of colors with respect to color detection and discrimination [10–13]. Anomalous trichromats and dichromats typically exhibit elevated thresholds along the red–green confusion axes in psychophysical tasks [14]. Neuroimaging studies further reveal a reduced sensitivity to reddish and greenish hues in early visual cortex (V1) in anomalous trichromats compared to trichromats, while in later visual areas (V2v & V3v) color contrast responses approached normal trichromat organization [15]. This finding indicates that post-receptoral compensation mechanisms may support suprathreshold color perception despite reduced cone signals [15–17]. Evidence from higher-level visual tasks also suggests substantial compensation beyond the photoreceptor level. Dichromats exhibit long-term memory for colored natural scenes that is identical to that of trichromats, including the same memory advantage for colored over achromatic images [18]. Together with neuroimaging studies, these findings suggest that higher-level representations of color may remain remarkably stable despite altered sensory input.

Assignments of single color samples to discrete color categories appear to be more preserved than might be expected. Bonnardel [19] examined and compared color categorization and naming in CVD and trichromat participants using free and constrained sorting of 140 Munsell samples and found that CVD participants’ category structures were broadly comparable to those of trichromats, despite differences in specific dimensions. Similarly, Cole et al. [20] reported that many mildly anomalous trichromats made few naming errors on a set of 10 basic colors, particularly when stimuli were large and saturated, although error patterns varied systematically across deficiency types.

Large color sample sets and naturalistic stimuli further support this pattern of relatively preserved color categories. Ma et al. [21] used a set of 208 Munsell chips to assess color constancy along a daylight axis and found that anomalous trichromats showed color categorization that was similar to trichromats across lighting conditions, whereas dichromats performed more poorly. Other work has shown that dichromats may nonetheless employ color terms in a manner that resembles typical naming patterns, particularly when non-chromatic cues (e.g., lightness or context) are available, and that there is considerable individual variability in naming performance among dichromats [22–26]. Uchikawa [22] further argued that this apparently normal color naming is likely supported by non-chromatic cues such as lightness, rather than by normal chromatic representations.

This suggests that categorical color behaviours found in individuals with red-green CVD are not simply a direct function of sensory loss, but may reflect preserved or learned categorical structures [27]. Despite the substantial literature on this topic, several questions remain open. First, most studies have emphasized group-level color naming patterns, whereas less is known about the reliability of category assignments over repeated measurements. Second, few studies have directly related categorical stability to quantitative measures of individual sensory precision. Third, it remains unclear whether color categories become progressively less stable as chromatic discrimination deteriorates from normal trichromacy through anomalous trichromacy to dichromacy.

In this study^1^, participants sorted 450 physical Munsell color chips into eleven labelled color categories and selected one representative chip for each category. This direct sorting task allowed participants to compare colors freely and to establish category boundaries using their own perceptual strategies. By repeating the task two hours later, we measured both agreement across observers and test-retest reliability within observers. Rather than focusing solely on discrimination accuracy or color naming, our approach quantifies the consistency, stability, and information content of categorical color assignments across trichromats, anomalous trichromats, and dichromats.

## Method

### Participants

The CVD group consisted of 20 male participants diagnosed using a combination of three assessments: a recent 24-plates edition of the Ishihara‘s tests [31] for color deficiency, the Nagel anomaloscope (Rayleigh match procedures) and the Color Assessment and Diagnosis (CAD) test [12]. The CAD test measures sensitivity thresholds for signal detection along the red-green and along the blue-yellow axis and provides quantitative measures as numeric standard units. We used the CAD RG scores as a measure for the severity of red-green deficits.

A participant’s deficiency subtype (deutan vs protan) was confirmed only when all three assessments agreed. Classification as anomalous trichromat or dichromat was based primarily on the anomaloscope results, as CAD severity scores may not distinguish these two categories reliably. The breakdown of the CVD group was 13 deuteranomalous, 2 protanomalous, 2 deuteranopes, and 3 protanopes.

The control group consisted of 10 participants (age range: 24-38 years, mean age: 30 years; 4 men, 6 women). Trichromatic vision was confirmed by using the 24-plates Ishihara test for color deficiency.

All participants provided written and informed consent. The study was approved by the Institutional Review Board at our department (Ethics approval # LEK 2020-0015) and conducted in accordance with the Declaration of Helsinki. Participants were reimbursed €20 per hour or received course credit for their time of participation.

### Setup

Testing was carried out under stable daytime illumination (Fig. 1). All participants were tested under comparable viewing conditions. Each participant was sitting at a table with eleven labelled containers corresponding to color category names in either English or German based on the participant’s preference (white, grey, black, red, orange, brown, yellow, green, blue, purple, pink). Stimuli were presented as a pile of physical randomly mixed Munsell color chips [32] in the center of table covered with a grey sheet of cloth.

**Figure 1.**
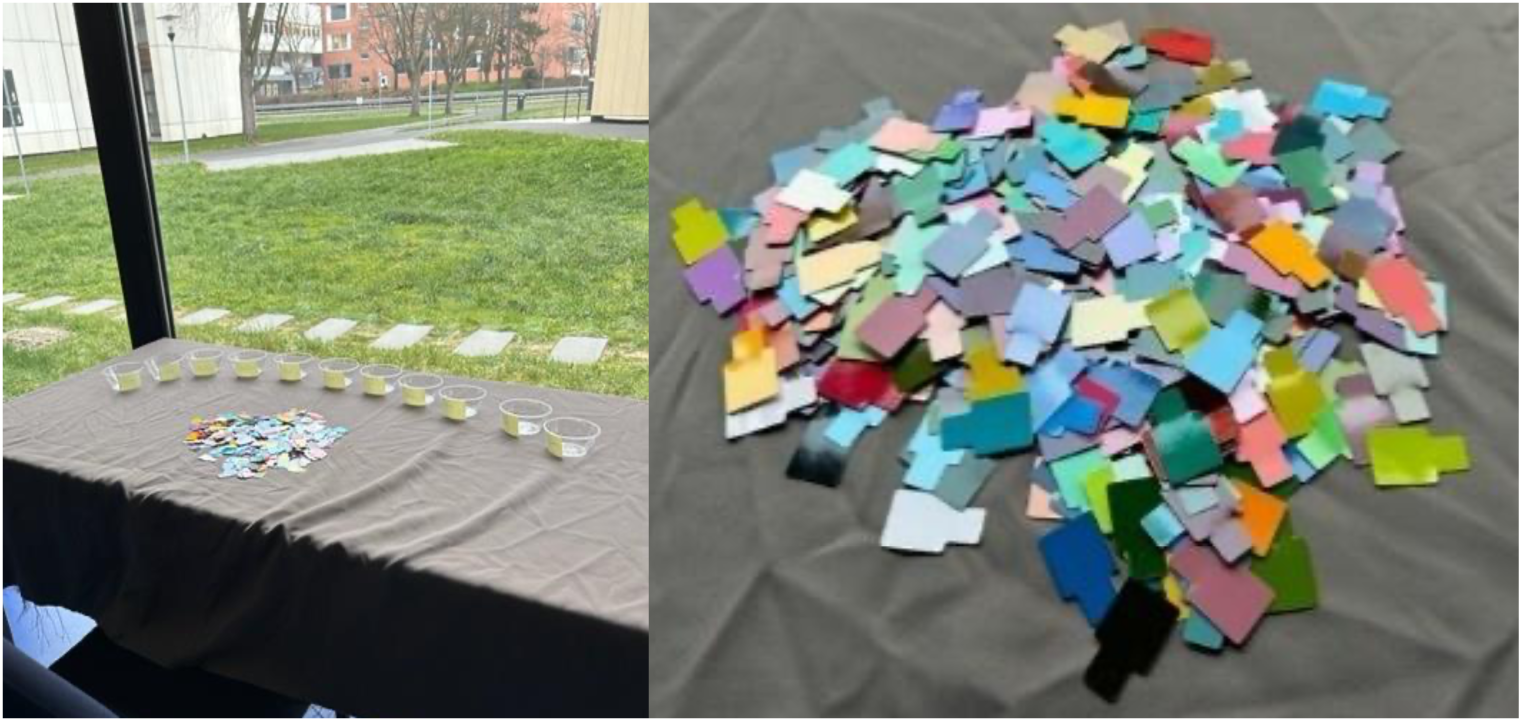
Experimental setup and stimulus presentation. Left: Participants sorted a pile of 450 physical Munsell color chips into eleven labelled containers corresponding to the basic color categories for which they also selected prototypical chips. The sorting task was performed on a grey background under natural daylight illumination. Right: The shuffled set of unsorted Munsell chips at the start of a sorting session.

### Stimuli

Stimuli consisted of 450 chips from the Munsell Glossy Edition, which included the 320 chips used in the World Color Survey (WCS) and additional 130 desaturated chips to ensure the coverage of low-chroma regions relevant for categorical assessment (Fig. 2). This construction of the Munsell chip set has been described previously in detail in the work of Olkkonen, Witzel, Hansen and Gegenfurtner [33], examining categorical color constancy with real surfaces. It is consistent with how WCS palettes have been defined in broader color categorization research [34].

**Figure 2.**
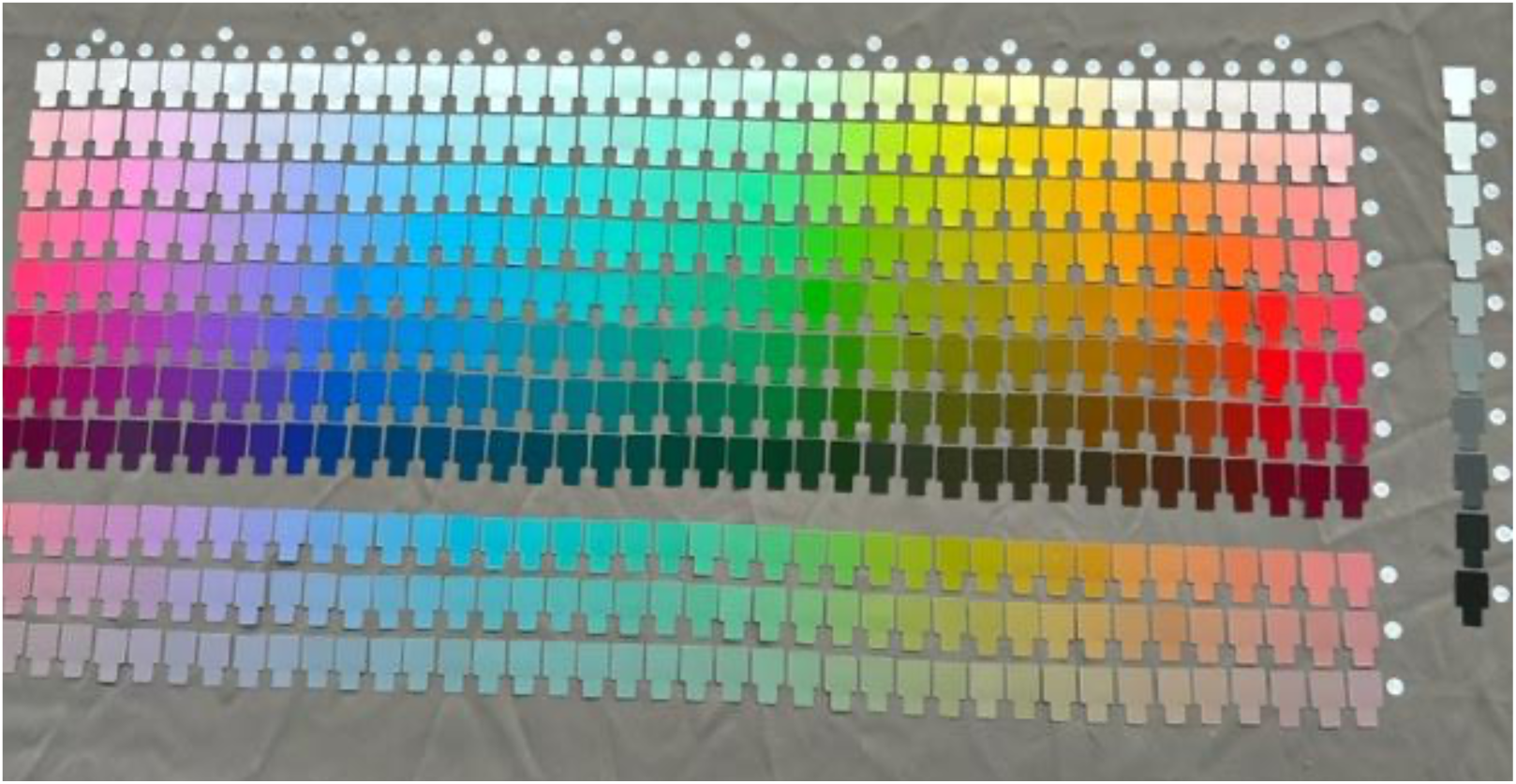
The sorted arrangement of chips. Munsell hue changes horizontally, while Munsell value increases from bottom to top. The lower three rows present chips of lower Munsell Chroma, the column to the right presents a series of achromatic chips.

### Design and Procedure

Participants contributing to the test-retest analysis completed two sorting sessions on the same day, separated by approximately two hours. This allowed the assessment of test-retest reliability. At the start of the first session (Time T1), participants signed consent forms and received verbal instructions. They were asked to sort a shuffled set of 450 Munsell color chips into the eleven labelled categories and to select one prototypical chip per category, namely the chip that best represented that category for them. Participants were instructed not to turn the chips over to inspect their backs. For control and questions one of the authors (AM) stayed in the room during the whole experiment. After a break of two hours, participants returned for the second part (Time T2) and repeated the sorting procedure with a newly shuffled set of chips. Sorting the chips typically required 45-60 minutes per session.

### Data Analysis

Preprocessing, statistical analysis and visualizations of the results were conducted in Python. To assess consistency of color categorization, we quantified agreements within groups (e.g., trichromats, anomalous trichromats, and dichromats) and test-retest reliability within individuals. For primary analyses, deuteranomalous and protanomalous observers were combined into a single anomalous trichromat group, and deuteranopes and protanopes were combined into a single dichromat group, consistent with previous studies [15,16], since both groups exhibit similar deficits along the red-green axis [35,36]. One dichromat participant did not complete T2 and was therefore excluded from any analysis involving test–retest.

Out of 450 chips, 217 showed complete agreement among all ten trichromatic participants (Fig. 5). These chips are referred to as consensus chips and served as a reference for subsequent analyses. To assess how representative each trichromat was of the group, we additionally used a leave-one-out procedure: for each trichromat in turn, the consensus name was recomputed from the remaining nine observers, and the held-out participant’s agreement with that nine participants’ consensus was recorded. This procedure provided an estimate of the variability expected among our small group of normal trichromats themselves.

#### Agreement analysis

To assess the reliability of categorical ratings across time points, we calculated pairwise Cohen’s κ [37] for all participant pairs within each color vision group (trichromats, dichromats, and anomalous trichromats) at T1 and T2. Cohen’s κ measures agreement corrected for chance and is defined as:

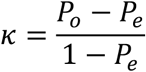

where *P_o_* is the observed proportion of agreement between raters, and *P_e_* is the expected proportion of agreement by chance. The expected agreement *P_e_* was estimated from the category marginals of the two raters. Agreement was also evaluated using permutation-corrected agreement and the Adjusted Rand Index (ARI) to confirm that conclusions were not dependent on the specific agreement metric or by differences in category marginals. Because pairwise agreement values are not statistically independent, group comparisons were carried out on one value per participant, namely each participant’s mean κ with the other members of their group. Test-retest reliability was calculated separately for each participant as Cohen’s κ between that participant’s T1 and T2 assignments, and these individual reliability values were compared across color vision groups using a Kruskal-Wallis test. Within-group agreement was also compared between sessions, using each participant’s mean κ with their own group at T1 and at T2.

#### Information-theoretic and color-space analyses

To characterise the sharpness of category boundaries independently of category frequencies, we computed for each chip the Shannon entropy (in bits) of the distribution of names assigned by the members of a group; entropy is zero when all members use the same name and increases as responses spread across categories. The correspondence between a group’s categories and the trichromat reference was quantified with the normalised mutual information between each member’s chip-by-chip names and the trichromat consensus name.

To locate where categories diverged in color space, each chip was represented by its Munsell hue, value, and chroma. We examined within-group agreement and agreement with the trichromat consensus as a function of Munsell hue across the full chip set. We also compared the Munsell value and chroma of chips on which a group’s majority name matched versus diverged from the trichromat consensus. Finally, reliance on lightness versus hue was indexed by the mutual information between the category label and binned Munsell value and, separately, binned Munsell hue, computed within each group. To estimate prototype stability, each chip was assigned its estimated CIELAB coordinates (L*, a*, b*) from the RIT Munsell renotation (adapted to a D65 white point), from which CIELAB DeltaE was calculated.

#### EnChroma condition

EnChroma lenses use spectral notch filtering intended to reduce overlap between L- and M-cone signals in anomalous trichromacy. To test whether this filtering altered color discrimination and categorical color sorting, a separate subset of nine anomalous trichromats completed an EnChroma condition. Participants first performed a baseline sorting without glasses and then repeated the task while wearing EnChroma glasses. The two sortings were separated by approximately two hours, during which participants wore the glasses continuously to allow adaptation before completing the second sorting.

Eight of the nine participants also completed a separate test-retest session on another day, in which they sorted the chips twice without glasses. The order of the EnChroma and test-retest days was counterbalanced across participants, whereas within the EnChroma session the baseline sorting always preceded the sorting with glasses. Differences between the baseline and EnChroma conditions were assessed using Wilcoxon signed-rank tests. Agreement within the EnChroma group and agreement with the trichromat consensus were quantified using the same procedures described above.

## Results

### Global Agreement Structure

To quantify agreements across participants, we computed pairwise Cohen’s κ values for color-category assignments across the full stimulus set. Within-group agreement was higher than between-group agreement (mean pairwise κ = 0.65 vs. 0.55). Because pairwise values are not independent, each participant contributing to many pairs, this was tested on one value per participant: each participant’s mean κ with their own group exceeded their mean κ with the other groups (0.64 vs. 0.56; Wilcoxon signed-rank W = 22, p < .001), indicating that participants with the same color-vision type showed more similar color sorting and naming behaviour than participants from the two other groups (Figure 3).

**Figure 3.**
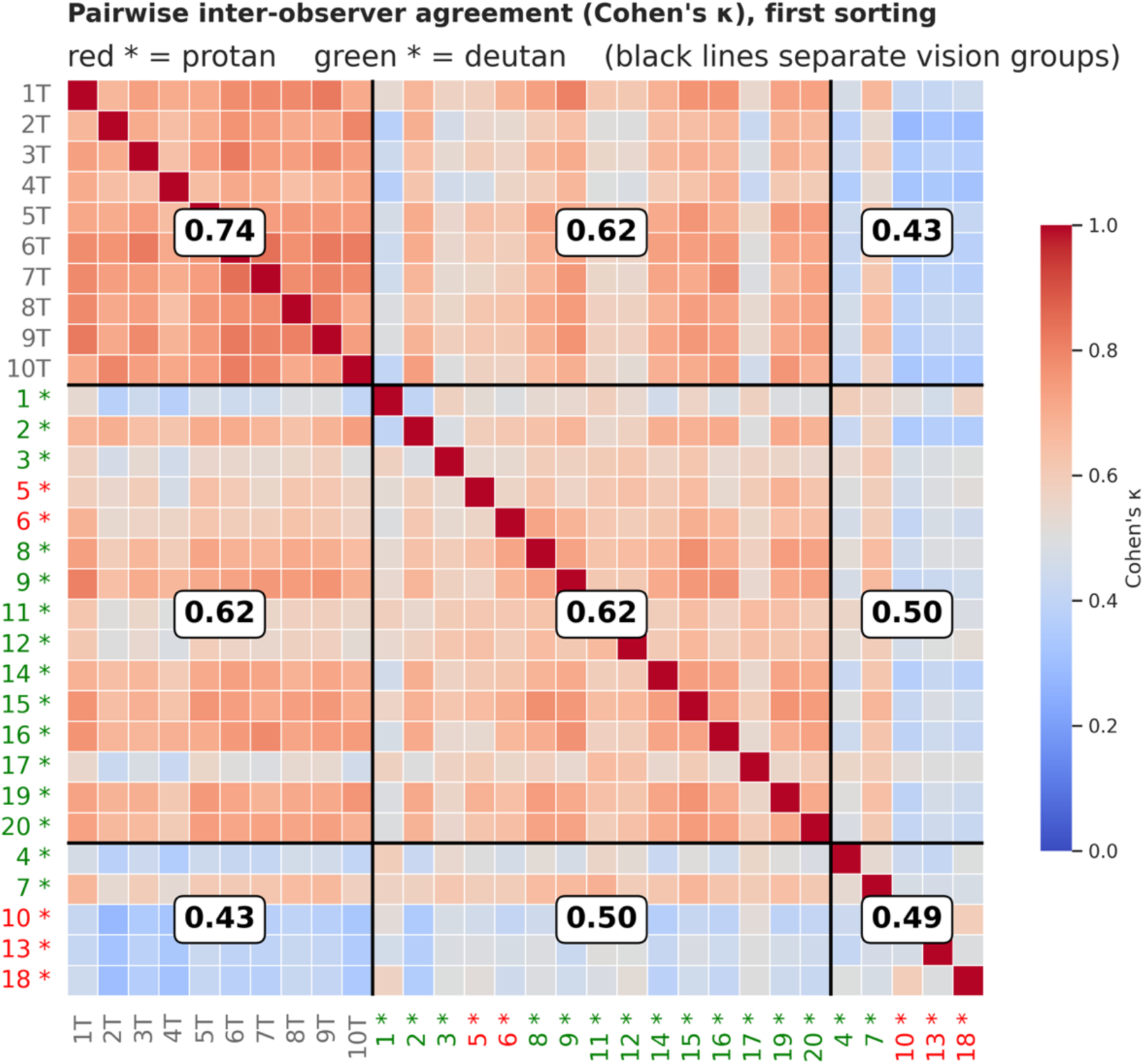
Matrix of pairwise inter-observer agreement. Each cell shows Cohen’s κ for the color-category assignments of two observers across the full set of chips. Observers are grouped by color-vision status. Stars mark protan and deutan subtypes within the anomalous trichromat and dichromat groups.

Agreement was graded by color vision status: highest among trichromats (mean within-group κ = 0.74), intermediate among anomalous trichromats (κ = 0.62), and lowest among dichromats (κ = 0.49; Figures 3 and 4). Equivalently, group members gave the same category name to 86%, 78% and 71% of chips on average. This ordering was statistically reliable. Computed on each participant’s mean within-group κ, the effect of group was significant (Kruskal–Wallis H = 23.87, p < .001), and Dunn tests with Holm correction separated all three groups (trichromats vs. anomalous trichromats p = .001; trichromats vs. dichromats p < .001; anomalous trichromats vs. dichromats p = .027).

**Figure 4.**
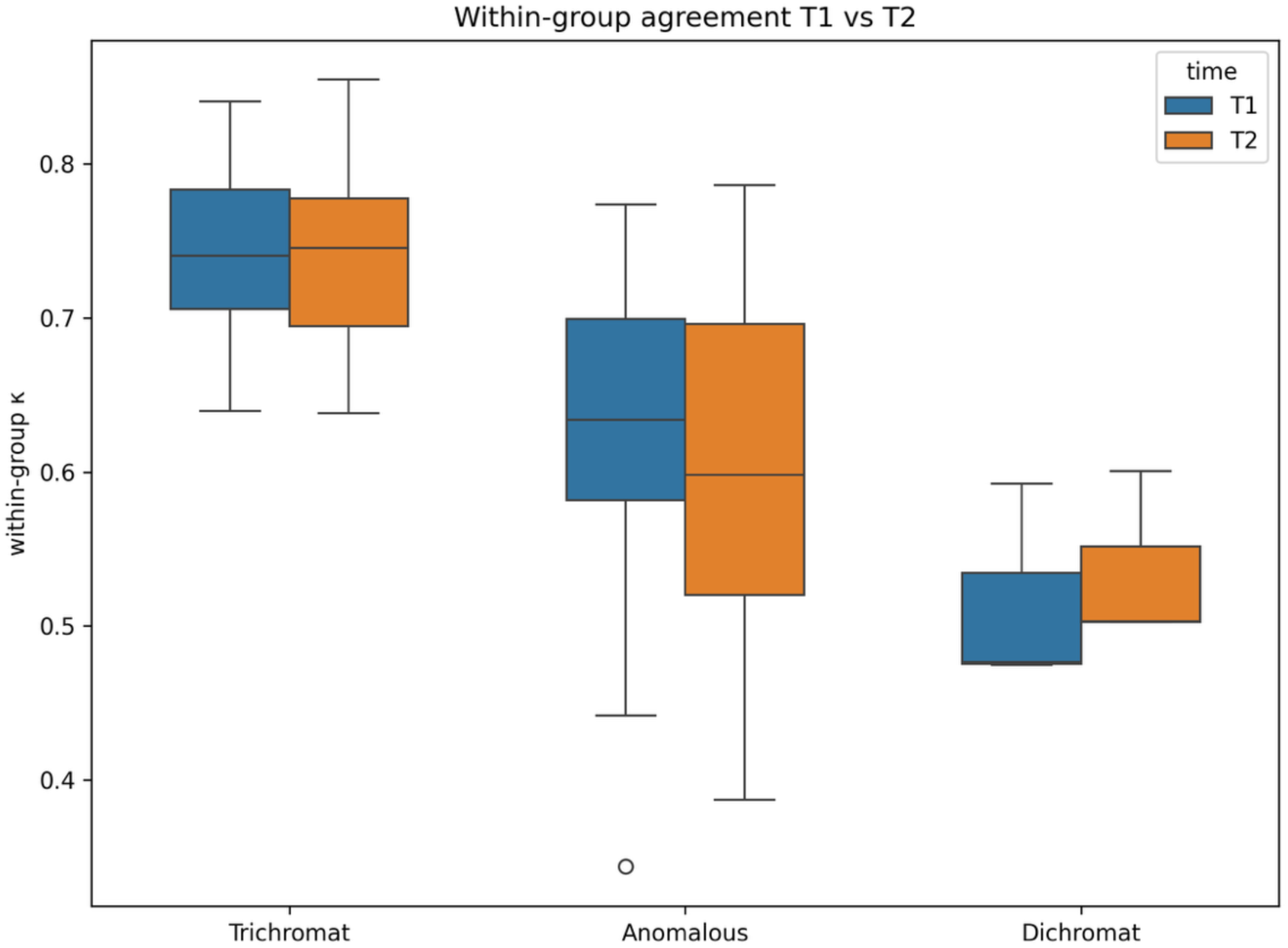
Within-group pairwise color category agreement. Agreement was quantified as mean Cohen’s κ between participants of the same color-vision group. Agreement was graded by color-vision status, being highest in trichromats, intermediate in anomalous trichromats, and lowest in dichromats. Agreement changed little between the two sessions.

We verified that this graded pattern was not driven by differences in category frequencies. Agreement above permutation expectation and the Adjusted Rand Index produced the same group ordering, with highest consistency among trichromats, intermediate consistency among anomalous trichromats, and lowest consistency among dichromats. Computed at the participant level, agreement above chance differed significantly across groups (Kruskal–Wallis H = 23.87, p < .001), as did the ARI (H = 23.22, p < .001). Thus, the main agreement results were robust across agreement metrics.

### Agreement with trichromat consensus

We next focused on the subset of chips for which trichromats showed unanimous categorical agreement. Across the full set of 450 chips, 217 were assigned the same color category by all ten trichromatic observers. These chips provided a conservative reference set for assessing how closely CVD observers followed trichromatic category patterns. Anomalous trichromats matched the trichromat consensus for 84.6% of these chips (range: 62–98%), indicating that their sorting of unambiguous surface colors to the eleven categories was close to that of trichromats for most of them, despite their measurable red-green discrimination deficits. Dichromats also showed substantial agreement with the categorical consensus, matching 60.6% of chips (range: 52–83%), but their sorting agreements were clearly reduced (see Figure 5). Thus, while anomalous trichromats used near-trichromatic categories for colors with unanimous trichromat labels, dichromats showed substantial but reduced agreements with the trichromat consensus. A leave-one-out analysis confirmed that the consensus was not driven by any single participant: each held-out trichromat agreed with the consensus of the remaining nine trichromats for 97.3% of chips on average (range: 91–100%).

**Figure 5.**
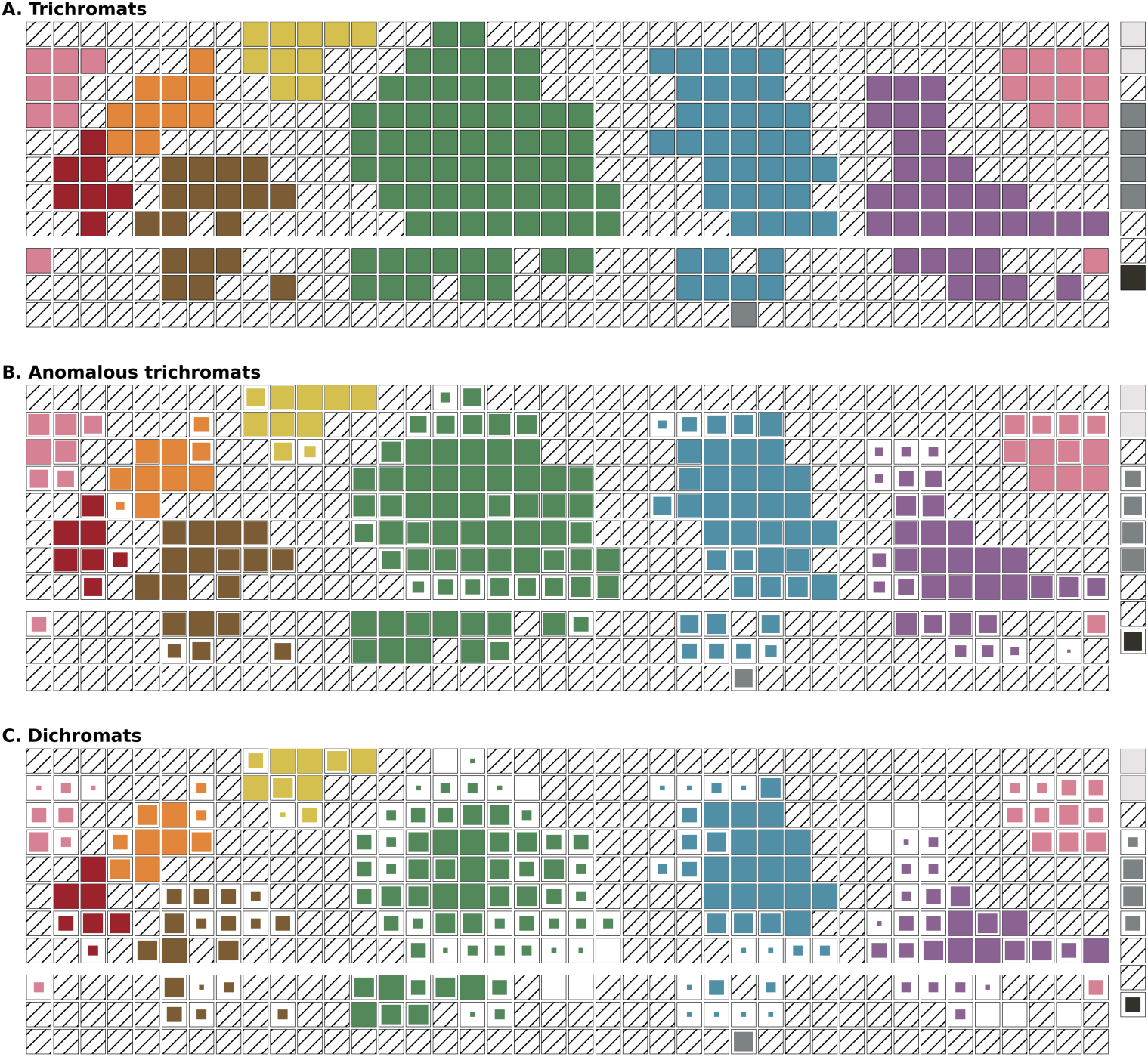
Consensus color chips of trichromats and the agreements with the two groups of color deficient participants. The top panel displays the 217 chips that all ten trichromats sorted into the same color categories, the trichromatic consensus. The middle panel shows the proportion of anomalous trichromats assigning each chip the trichromat consensus name, and the bottom panel shows the corresponding proportion for dichromats. For each chip, the color of the square indicates the consensus color name, and the size of the square indicates the degree of agreement with that name. Larger squares indicate higher agreement. Anomalous trichromats closely followed the trichromat consensus, whereas dichromats showed lower but still substantial agreement.

A complementary question is how each group’s internal consistency depended on chip ambiguity. We compared within-group agreement, the proportion of a group’s members assigning the modal category, separately for the 217 unanimous (consensus) chips and the remaining 233 nonconsensus chips. For trichromats, agreement fell from 100% on the consensus chips (which are defined by unanimous trichromat labelling) to 72% on the nonconsensus chips. Anomalous trichromats showed a similar but smaller decline, from 85% to 71%. Dichromats, by contrast, were essentially as consistent on the ambiguous chips as on the unanimous ones, at 72% and 70% respectively. As a consequence, the three groups were separated by 28 percentage points on the consensus chips but converged to within two points of one another (70 to 72%) on the nonconsensus chips. The graded group differences in category consistency were thus carried almost entirely by the unambiguous surface colors. For the chips that even trichromats found hard to categorise, dichromats were no less consistent than trichromats. Because the groups differ in size and the modal proportion depends on sample size, we repeated the comparison with group size equalised at five observers by repeatedly subsampling the trichromat and anomalous trichromat groups. The convergence persisted: on the non-consensus chips the matched-size estimates were 74%, 74% and 70% for trichromats, anomalous trichromats and dichromats, whereas the difference on the consensus chips remained large (100%, 86% and 72%).

### Test-Retest Reliability

The high agreement with trichromat consensus chips shows that many categorical assignments were preserved in CVD, especially in anomalous trichromats. This raised the question of whether these assignments were also stable within individual participants across repeated measurements. Test–retest reliability was quantified for each participant as Cohen’s κ between their own T1 and T2 sortings of the full chip set, and these within-participant values were compared across the three groups. Reliability was highest in trichromats (mean κ = 0.86), lower in anomalous trichromats (mean κ = 0.74), and lowest in dichromats (mean κ = 0.70); the difference across groups was significant (Kruskal–Wallis H = 15.2, *p* < .001). Although all three groups re-used their color names fairly consistently across the two-hour interval, categorization became progressively less reliable with the severity of the color vision deficiency, with dichromats the least stable (Figure 6).

**Figure 6.**
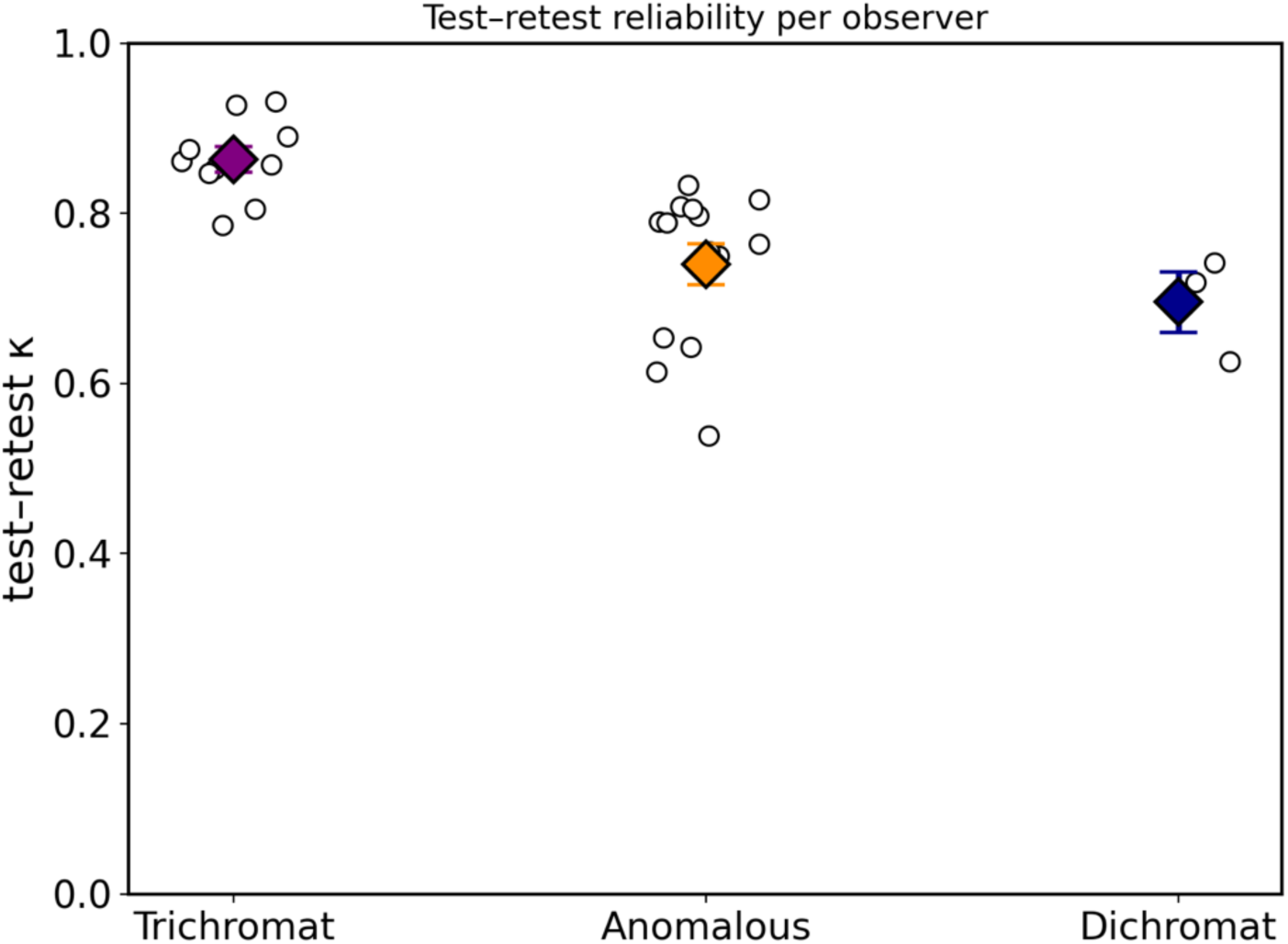
Test-retest reliability of color categorization. Within-participant agreement between the two sessions was quantified as Cohen’s κ between each participant’s T1 and T2 sortings across the full chip set. Points show individual participants and boxes summarize each color-vision group.

### Category Consistency and Sensory Precision

The high agreement of anomalous trichromats with the trichromat consensus raises the question of whether the remaining variability in categorization is related to the degree of sensory loss. We therefore examined whether individual CAD scores predicted color-category consistency. Higher RG-CAD scores of anomalous trichromats (n = 15), indicating poorer red-green discrimination, were associated with lower category consistency, measured as the mean pairwise Cohen’s κ with other anomalous trichromats at the first sort (Spearman r = −.704, *p* = .003). Thus, even though anomalous trichromats showed near-trichromatic categorization for consensus chips, individuals with more severe sensory losses showed less consistent categorical assignments. In dichromats (n = 5), no reliable association was found (Spearman r = −.100, *p* = .873), although this analysis was limited by the small sample size. These results indicate that color categories in anomalous trichromats are largely preserved, but the fidelity of category assignment is still constrained by residual sensory precision (Figure 7).

**Figure 7.**
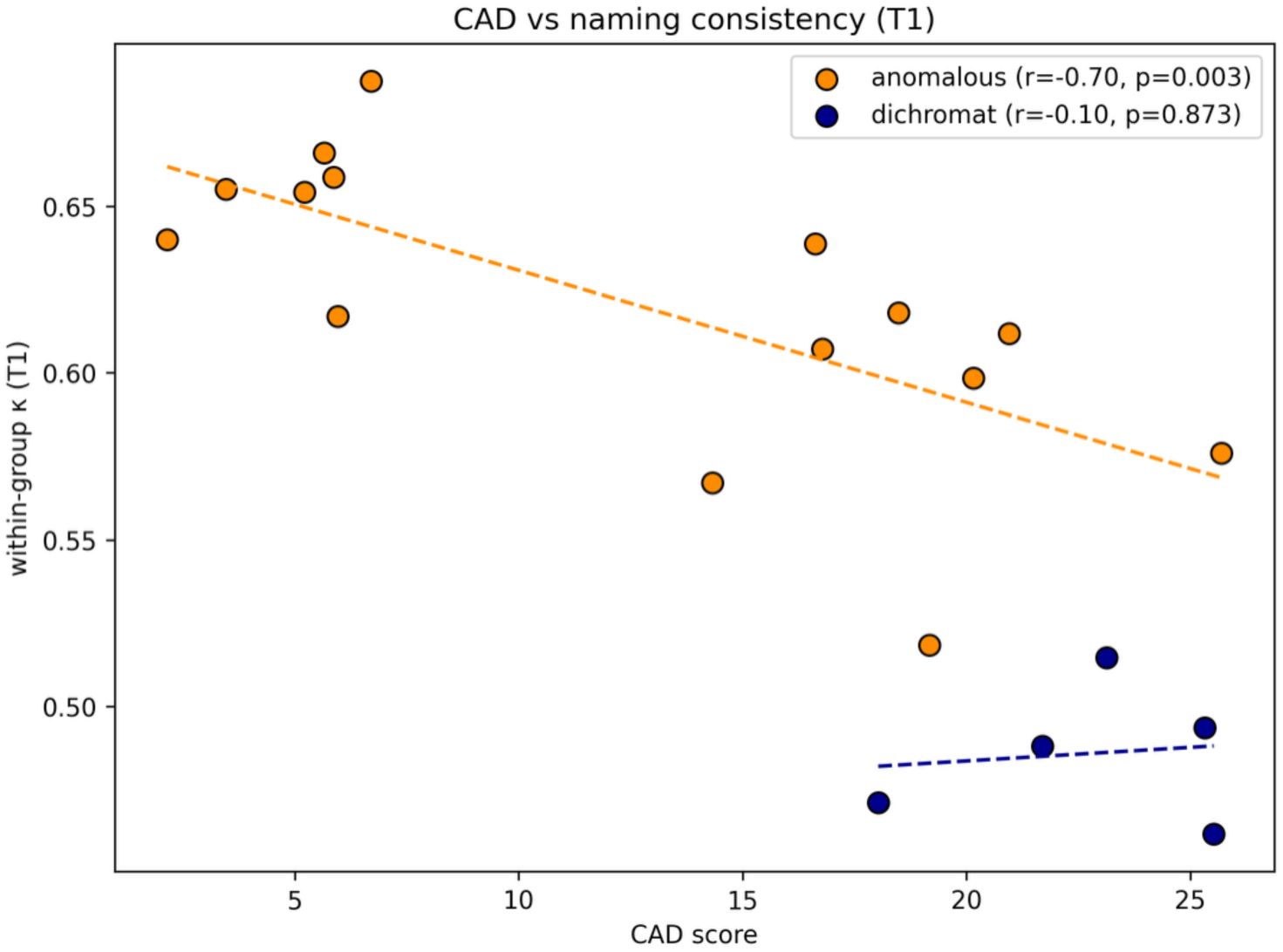
RG-CAD scores and color-category consistency. For each color-deficient participant, his CAD red-green discrimination threshold (x-axis) is plotted against his category consistency measured as the mean within-group pairwise Cohen’s κ at the first sort. Higher RG-CAD scores indicate higher red-green discrimination thresholds. Among anomalous trichromats, poorer red-green discrimination was associated with lower category consistency. No reliable relationship was found for our small group of dichromats. Dashed lines show separate linear fits for each group.

### Distribution of color-category use

The preceding analyses quantified color category agreements between participants and across repeated measurements. We also examined whether the groups differed in how often they used each color category for chip assignments. Because the category-use distributions were nearly identical at T1 and T2, responses were combined across sessions for visualization. Trichromats and anomalous trichromats showed similar distributions, with Green and Blue among the most frequently used labels in both groups. Dichromats showed a more even distribution across categories and used Grey and White more frequently than the other groups. Thus, category-use distributions were stable across sessions but differed systematically across color-vision groups, with dichromats using achromatic labels more frequently (Figure 8).

**Figure 8.**
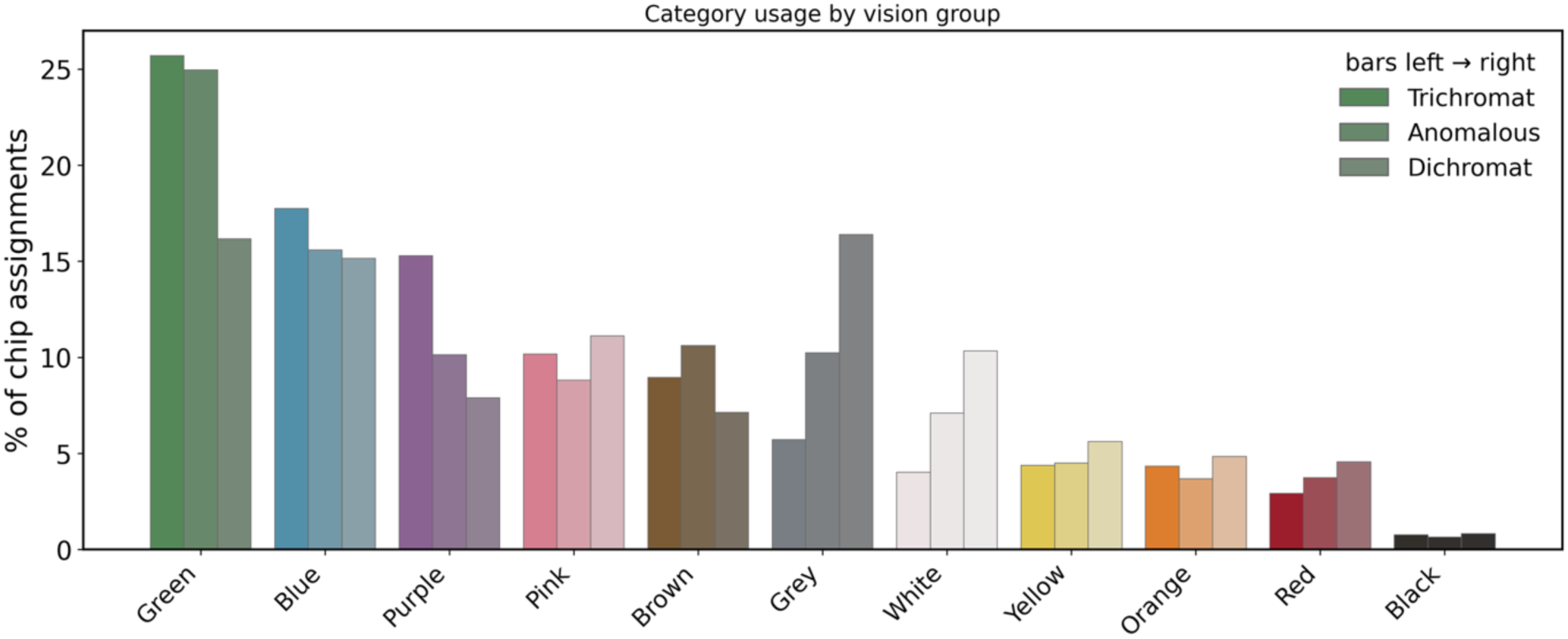
Distribution of color-category use across groups. Proportion of chip assignments to each of the 11 color categories, pooled across T1 and T2. T1 data are included for all observers, and T2 data for the subset who completed the retest. Categories are ordered by overall frequency. Within each category, bars show trichromats, anomalous trichromats, and dichromats from left to right. Bar colors indicate the corresponding category label, with decreasing saturations used to distinguish the three groups of participants. Trichromats and anomalous trichromats used similar assignments to the different color categories, whereas dichromats used the achromatic categories of Grey and White more frequently and distributed responses more evenly across categories.

### Prototypical Color Selection and Stability Over Time

Beyond assigning all chips to color categories, participants also selected one prototypical chip for each category. This allowed us to ask whether the internal reference points for the color categories were stable across the two-hour interval. For each observer and category, we compared the prototypical chip selected at T1 with the prototypical chip selected at T2. Stability was quantified in two ways: first, by asking whether the prototype selected at T1 still belonged to the same category at T2, and second, by measuring the color-space distance between the T1 and T2 prototypes.

Prototype stability differed across color-vision groups. Trichromats were highly stable, with all T1 prototypes remaining within the same category at T2. Anomalous trichromats also showed high stability with 86.36% of prototypes remaining in the same category, whereas dichromats were less stable with 43.18% of prototypes remaining in the same category. These group differences were highly significant (Kruskal-Wallis H = 20.07, *p* < .001). Posthoc pairwise comparisons showed that trichromats differed from both anomalous trichromats (*p* = .002) and dichromats (*p* < .001), and that anomalous trichromats were more stable than dichromats (*p* = .049).

A complementary analysis measured the distance between the T1 and T2 prototype selections directly in color space. The mean CIELAB shift increased from trichromats (mean ΔE = 7.7) through anomalous trichromats (12.5) to dichromats (18.0), and this group difference was significant (Kruskal-Wallis H = 8.9, *p* = .011; Figure 9). The prototype arrays in Figure 10 provide a complementary view of this pattern, showing tightly clustered prototype selections in trichromats and anomalous trichromats and substantially greater dispersion in dichromats. Thus, CVD affected not only category assignment but also the stability of category prototypes, with dichromats showing the largest shifts in their selections across repeated measurements.

**Figure 9.**
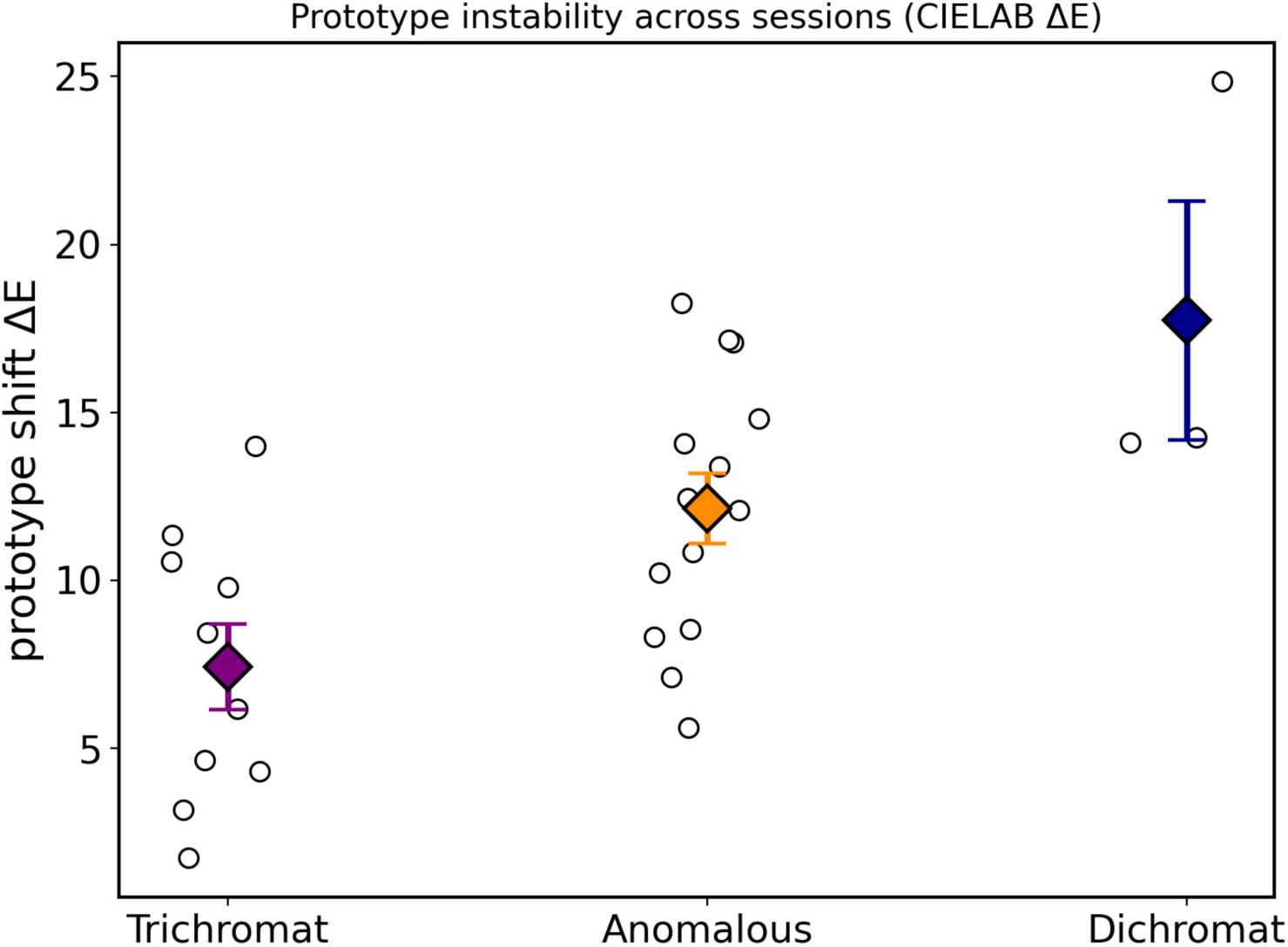
Prototype instability across sessions. For each observer and color category, the prototypical chip selected at T1 was compared with the prototypical chip selected at T2 two hours later. Their separation was quantified as the CIELAB color difference ΔE. Larger values indicate less stable prototype selection. Each point shows one observer’s mean ΔE across categories and boxes summarize each color-vision group.

**Figure 10.**
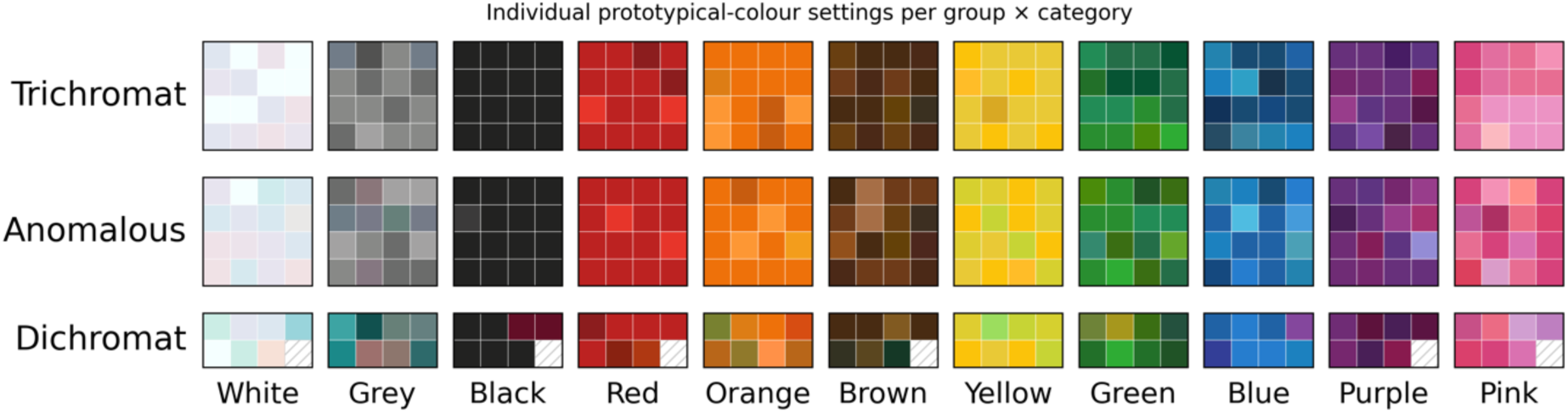
Individual prototype selections by color vision groups and color categories. Each block shows the actual chips chosen as the best example ("prototype") for a color category as single patch selections pooled across sessions. Rows correspond to the three groups and columns to the eleven categories. For trichromats and anomalous trichromats a random sample of 16 selections is shown per cell; for dichromats all selections are shown, with hatched patches marking missing settings.

### Color-Space Structure, Confusions, and Information Content

To localise where color sorting agreements broke down, we examined within-group agreement and agreement with the trichromat consensus as a function of Munsell hue across the full chip set (Figure 11). Color assignments of anomalous trichromats closely followed trichromats across hues, whereas the agreements of dichromats were strongly hue-dependent. Agreements with the trichromat consensus were lowest across green to blue-green hues (Munsell steps approximately 17 to 28), where neighbouring chips are separated mainly by the L minus M signal that is unavailable or strongly degraded in red-green CVD. A secondary dip occurred in the purple region, whereas agreement remained relatively high for reds and blues. Averaged over hue, dichromats matched the trichromat consensus on 51% of chips, whereas anomalous trichromats matched it on 70%. Within-group agreement showed the same ordering, being lower in dichromats (0.71) than in anomalous trichromats (0.78) or trichromats (0.85). Agreement was also reduced for low-chroma chips in all three groups, consistent with weaker chromatic signals producing less stable categories.

**Figure 11.**
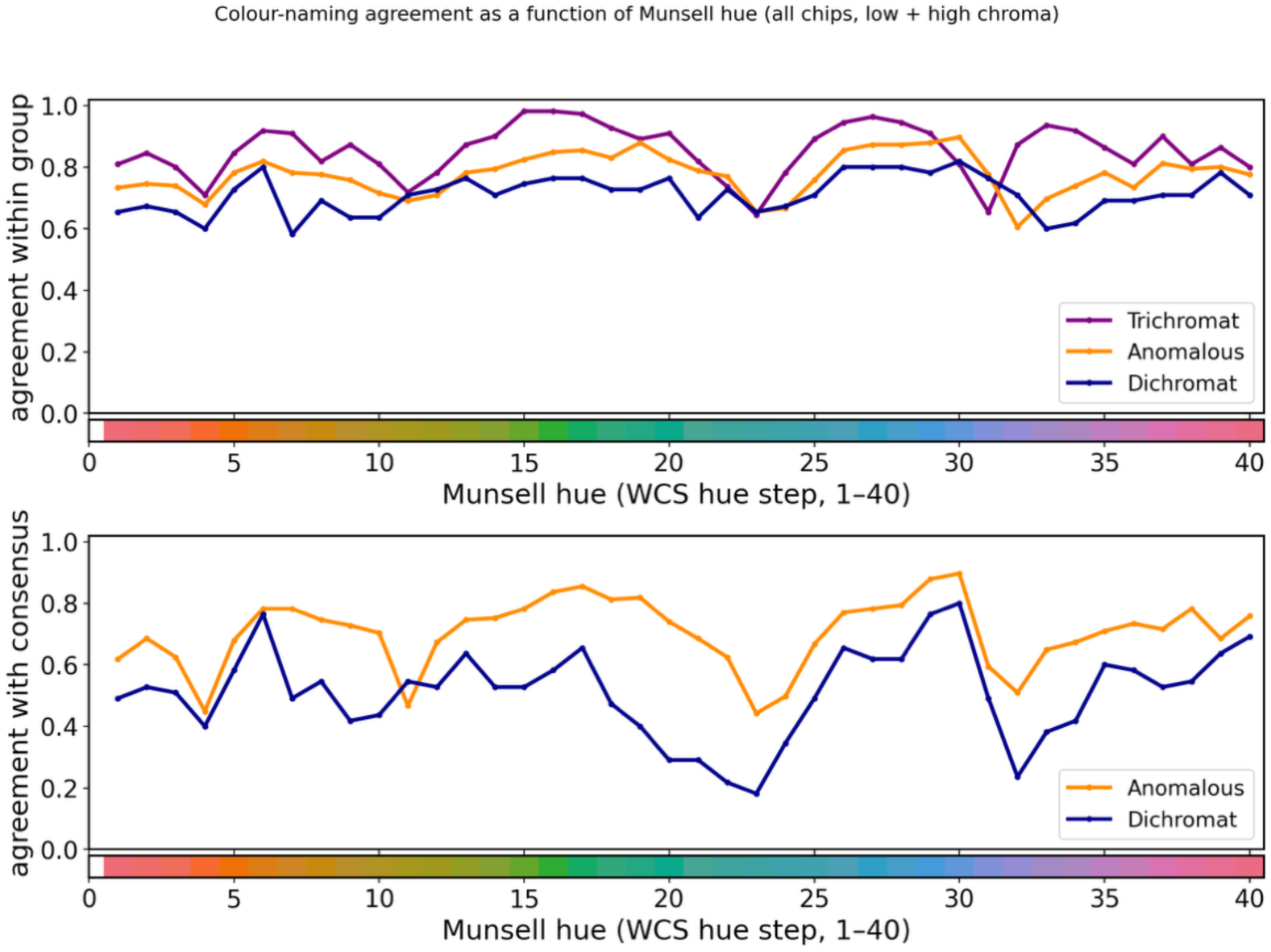
Color-category agreement as a function of Munsell hue. Agreement is shown across the 40 WCS hue steps for the full chip set. The top panel shows within-group agreement for trichromats, anomalous trichromats, and dichromats. The bottom panel shows agreement with the trichromat consensus for anomalous trichromats and dichromats. Dichromat agreement falls most strongly across green to blue-green hues, with a secondary reduction in the purple region, and remains higher for reds and blues. The colored strip below each panel marks the corresponding chip hues.

To examine the categorical structure of these differences in color assignments, we cross-tabulated the color category chosen for each chip against the trichromat consensus by pooling T1 and T2 because their matrices were nearly identical across sessions. Agreement with the trichromat consensus, corresponding to the matrix diagonal, decreased from trichromats (83%) through anomalous trichromats (71%) to dichromats (59%). The off-diagonal responses were systematic rather than random. Dichromats most often interchanged green, yellow, and brown, or assigned these chips to grey. They also frequently named purple chips blue and near-black chips brown. These confusions are consistent with reduced information along the red-green dimension and increased reliance on residual chromatic and lightness cues. Anomalous trichromats showed a similar response pattern in much weaker form (Figure 12). Overall, anomalous trichromats and dichromats assigned the same majority category to 73% of chips, indicating that the two CVD groups share much of their category map even though they differ in internal consistency and in how individual categories are applied.

**Figure 12.**
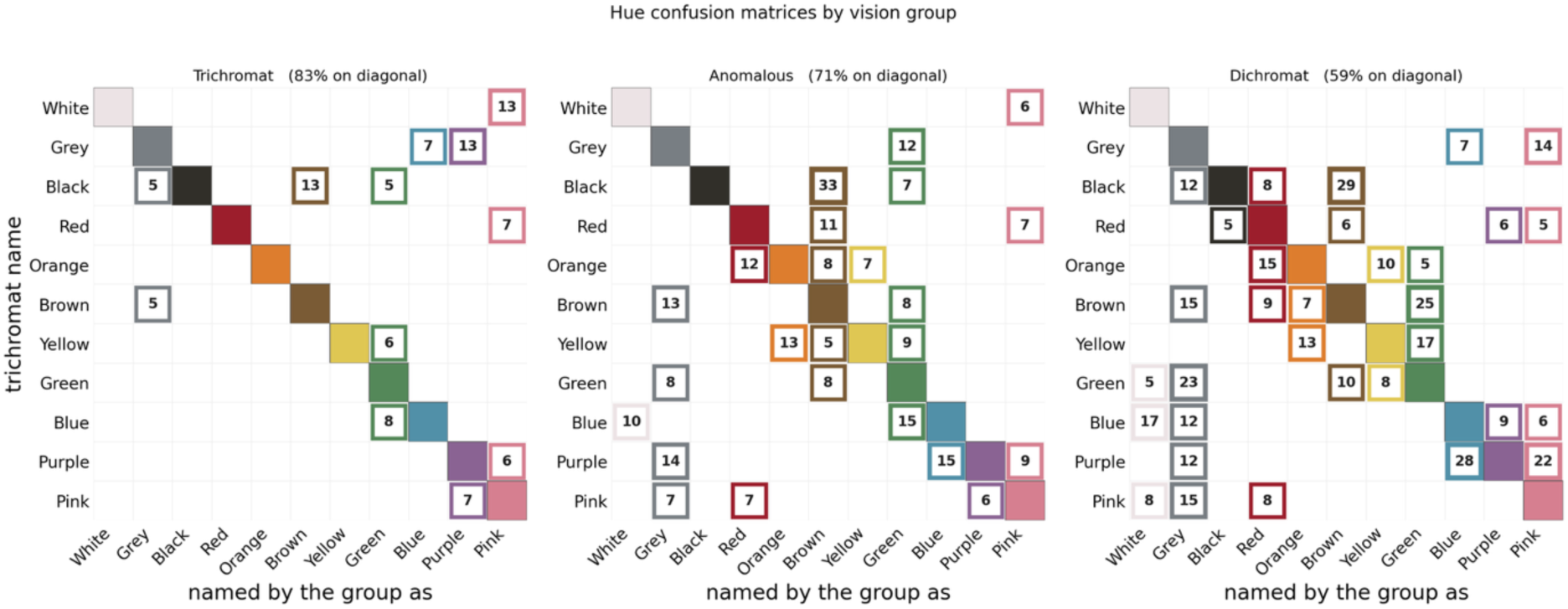
Color-name confusion matrices for the three groups tested. The color-vision group is given above each color-name confusion matrix, rows show the trichromat consensus color name for each chip and columns the names assigned by members of each group. Data are pooled across T1 and T2, and each row is normalised to 100%. Diagonal cells display the agreement with the trichromat consensus and are filled with the corresponding category color. Off-diagonal confusions of at least 5% are framed in the color assigned by the group, with the row percentage shown. Trichromats are largely diagonal, whereas dichromats show systematic confusions involving green, yellow, brown, grey, purple, blue, and near-black chips. Anomalous trichromats show a similar but weaker pattern.

We then quantified the sharpness of category boundaries with Shannon entropy. For each chip, the distribution of names assigned by a group was treated as a probability distribution, and entropy was computed in bits. Entropy is zero when all observers use the same name and increases as responses are distributed across multiple categories. Mean per-chip entropy increased from trichromats (0.46 bits) to anomalous trichromats (0.76 bits) to dichromats (0.84 bits; Kruskal-Wallis H = 104.9, *p* < .001). Thus, sensory loss did not eliminate categorical structure, but made category assignments less sharply defined. In parallel, normalised mutual information between a group’s chip-by-chip names and the trichromat consensus name declined from trichromats (0.72) to anomalous trichromats (0.54) to dichromats (0.41; Figure 13). This indicates that the trichromat consensus category became progressively less predictive of the names assigned by observers with stronger color vision deficiency.

**Figure 13.**
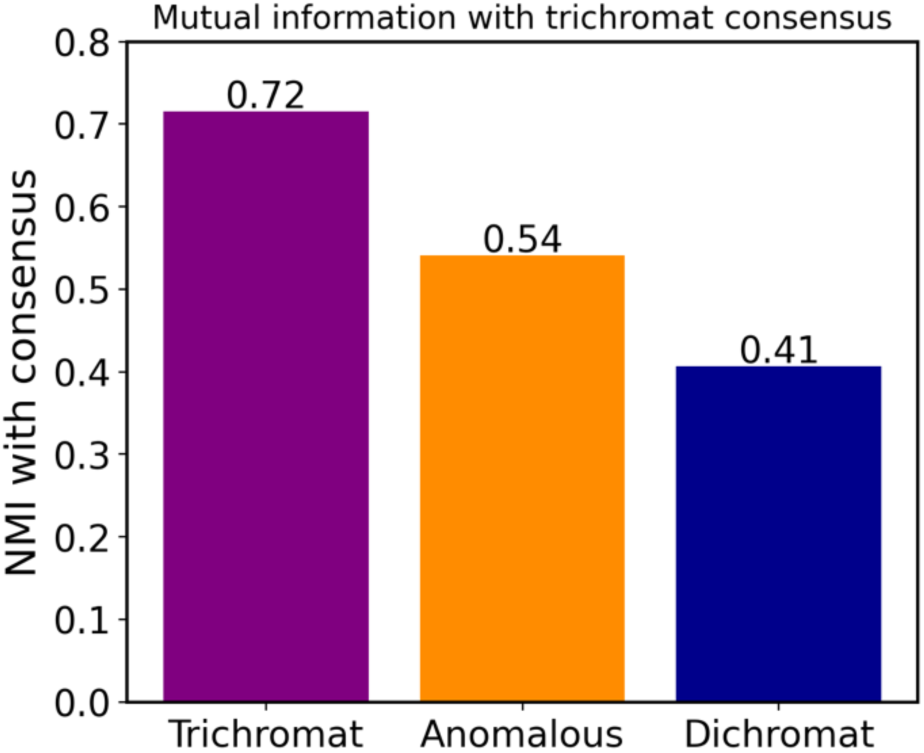
Normalised mutual information between group categories and the trichromat consensus. Higher values indicate that the group’s chip-by-chip color names are more strongly determined by the trichromat consensus category. Mutual information decreased from trichromats (0.72) through anomalous trichromats (0.54) to dichromats (0.41).

Finally, we tested whether color-deficient observers relied more strongly on lightness when sorting the hues of chips. For each group, we computed the mutual information between category labels and discrete Munsell value, and separately between category labels and discrete Munsell hue. Munsell hue carried more information than value in every group, but the ratio of value information to hue information increased monotonically from trichromats (0.25) through anomalous trichromats (0.49) to dichromats (0.70). This pattern indicates progressively greater reliance on lightness as chromatic signals weaken (Figure 14).

**Figure 14.**
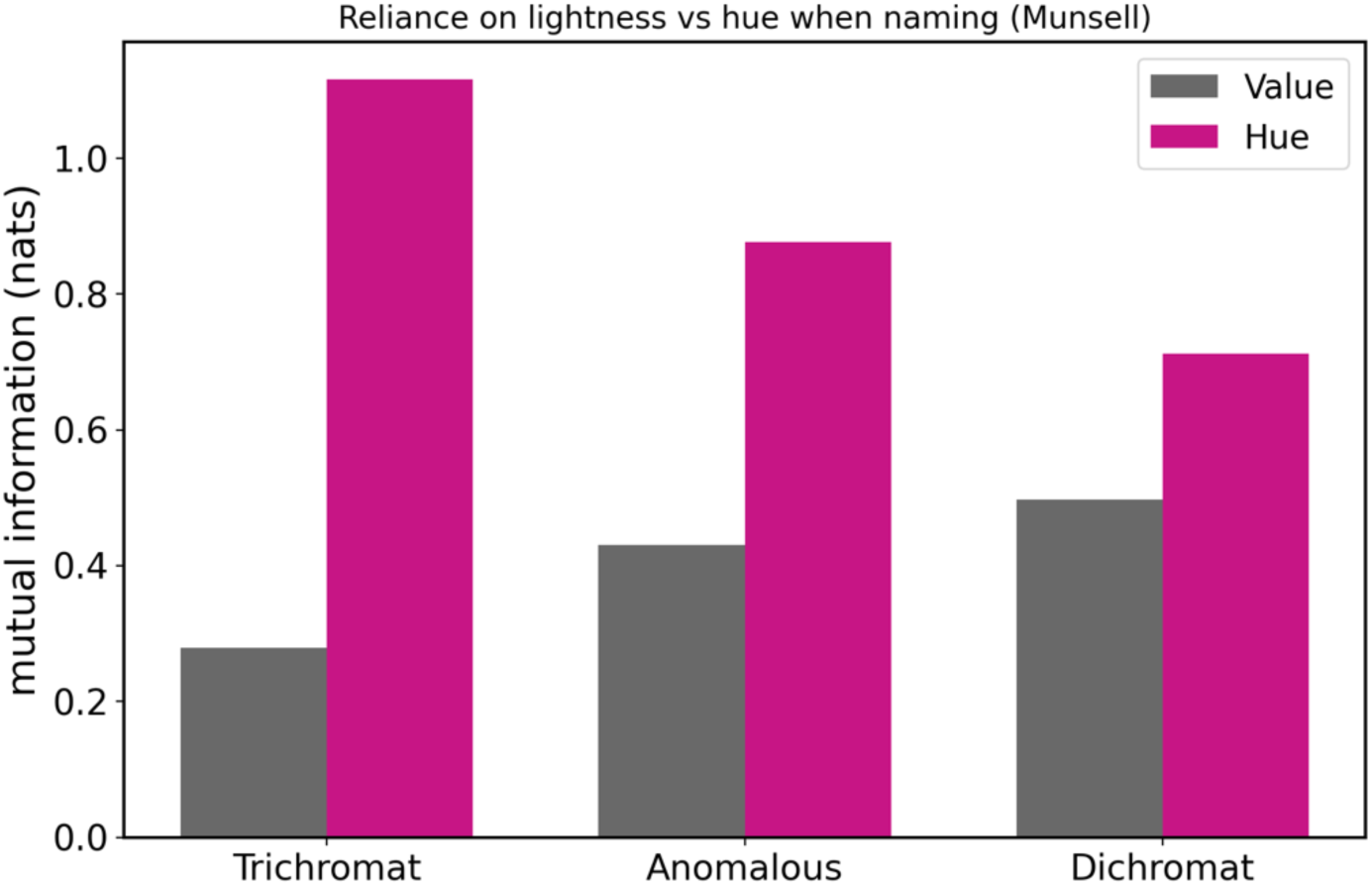
Mutual information between color-category labels and Munsell value or Munsell hue. For each group, mutual information was computed separately for binned Munsell value and binned Munsell hue. The value-to-hue ratio, annotated above each pair, increased from trichromats to anomalous trichromats to dichromats, indicating progressively greater reliance on lightness as chromatic signals weaken.

### Enchroma

We also tested whether acute spectral filtering with EnChroma lenses would make colour sorting and categorization in anomalous trichromats more similar to the trichromat consensus. Nine anomalous trichromats completed the EnChroma condition after approximately two hours of continuous adaptation. Eight participants completed both the EnChroma and separate test-retest sessions and therefore provided four chip sortings across two days; one participant completed only the EnChroma session.

Overall color sorting consistency did not change (Figure 15). Mean pairwise Cohen’s κ within the EnChroma group was virtually identical without glasses (κ = .59) and with glasses (κ = .58; Δκ = −.01, Wilcoxon p = .539). Agreement with the trichromat consensus for the 217 consensus chips increased only slightly, by .02 on average, but this change was not significant (Wilcoxon p = .24; Figure 15). Category-use distributions were also similar across the two conditions. Green and blue were among the most frequent labels both without glasses and with glasses, and there was no systematic shift toward the trichromat reference pattern. These participants had also completed a separate test-retest session without glasses, so agreement with the trichromat consensus could be followed across all four sortings; it stayed within the same range throughout, indicating that the small differences between conditions reflected ordinary test-retest variability rather than an effect of the glasses on color perception.

**Figure 15.**
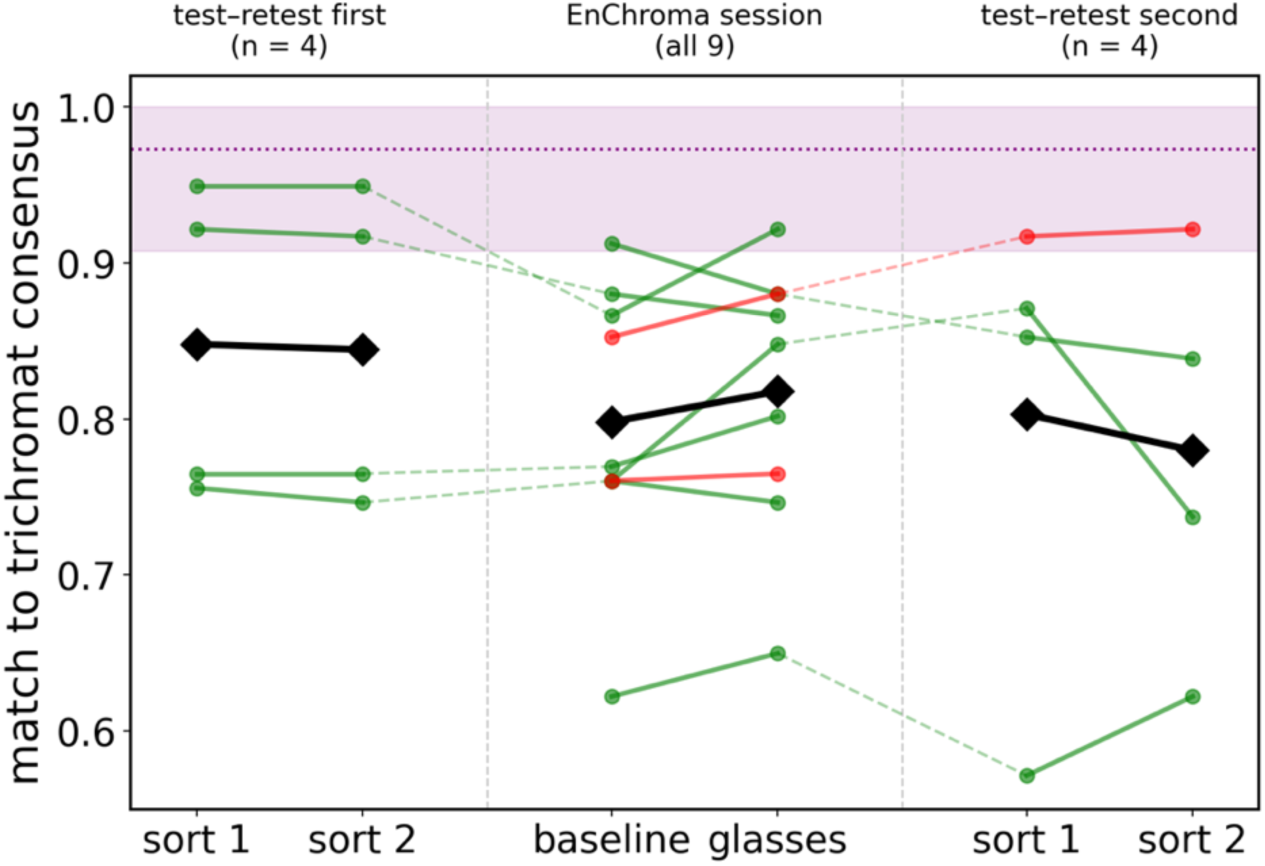
EnChroma condition across all color sortings. One group of four participants completed a test-retest session with two sortings without glasses followed by an EnChroma session with a baseline sorting without glasses and a sorting while wearing the glasses, the other group of four started with the EnChroma session so that the order of the two sessions was counterbalanced. One participant completed only the EnChroma session. The y-axis shows agreement with the trichromat consensus, the proportion of the 217 consensus chips. Colored lines connect each participant across their sortings (green: seven deuteranomalous; red: two protanomalous), with solid segments within sessions and dashed segments bridging the two days; black diamonds are group means within each block. The shaded band and dotted line displays the trichromat reference obtained by leave-one-out, in which each trichromat was compared with the consensus of the other nine.

Thus, wearing EnChroma glasses for approximately two hours and while completing the sorting did not produce a measurable change in color categories in anomalous trichromats. The result does not rule out longer-term adaptation, but it indicates that the lenses did not produce a large immediate shift toward trichromatic category structure.

## General discussion

In a standard dichromat simulation, the Munsell array appears reduced mainly to yellow, blue, white, grey, and black (Figure 16). Such simulations are useful for illustrating how severely chromatic signals are compressed, but they should not be taken as literal depictions of dichromat experience. Broackes [38], drawing on cases of unilateral color vision deficiency, argued against the view that dichromatic experience is exhausted by two hues, and suggested that it may be considerably richer and more context-dependent than such reductions imply (see also [3]). Against this background, the present results are striking: categorical color structures were not lost in CVD participants. Instead, color categories persisted, but their stability and sharpness declined with the severity of sensory loss. Anomalous trichromats matched the trichromat consensus for 85% of unanimously named chips and and retained substantial test–retest reliability, though reliably below that of trichromats. Dichromats showed lower agreement and greater prototype instability, but their responses remained structured rather than random. The principal effect of CVD was therefore not a collapse of color categories, but a graded loss of precision in assigning colors to otherwise recognizable categories.

**Figure 16.**
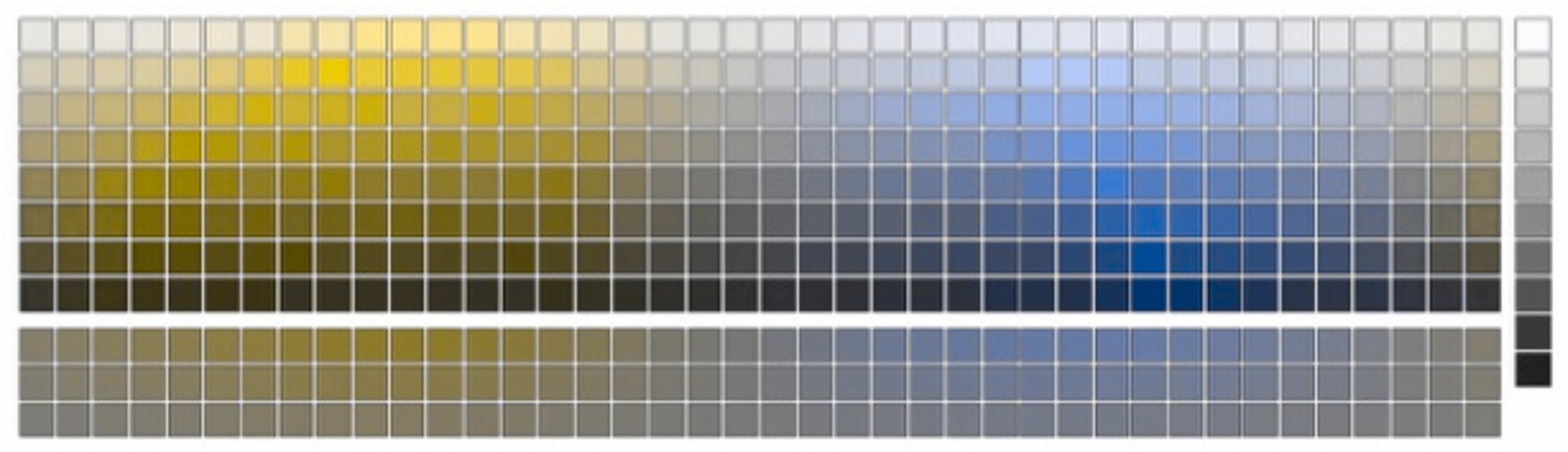
Simulated deuteranopic appearance of the Munsell chip set. The same Munsell chip arrangement shown in Figure 2 is displayed after simulation of deuteranopic vision using the algorithm of Brettel, Viénot and Mollon [39], and implemented with the software provided by DaltonLens. The simulation illustrates the severe compression of chromatic appearance, especially the apparent reduction to yellow, blue, white, grey, and black. The simulation is illustrative only and was not used in the statistical analyses.

### Relationship to earlier studies

These findings align with those reported in earlier studies showing that color categorization and naming can remain relatively preserved despite well documented sensory deficits along the red-green axis [19,20,22,23,27,40–42]. Our results extend these findings in two ways. First, they show that anomalous trichromats can preserve highly trichromat-like category structures across a dense sampling of physical Munsell chips, including low-chroma regions in which sorting and naming might be expected to be unstable. Second, they show that dichromat sorting is not random or collapsed into a small set of labels, but remains systematically organized despite reduced agreements with trichromats. The high correspondence between the chip sorting and categorization of anomalous trichromats and trichromats suggests that altered red or green cone sensitivities do not necessarily lead to a large-scale reorganization of color category structure.

The dichromat results further clarify how color-category structure changes when chromatic information is severely reduced. Dichromats differed more strongly from trichromats, however their responses nevertheless showed coherent category structures and substantial within-group agreements. The information-theoretic analyses made this pattern concrete. The share of category information carried by lightness relative to hue rose monotonically across groups, with value-to-hue mutual-information ratios of 0.25, 0.49, and 0.70 for trichromats, anomalous trichromats, and dichromats, respectively. This pattern is consistent with progressively greater reliance on lightness as chromatic signals weaken. It also fits earlier proposals that dichromats can use non-chromatic cues, especially lightness, to assign color names to surfaces that are difficult to distinguish chromatically [22,23,27,40–42].

How are color categories preserved despite degraded sensory input? The results of the present study suggest that categorical color sorting is supported by stable learned category templates, but that the reliability with which individual chips can be assigned to those templates depends on the available sensory evidence. In anomalous trichromats, this dependence was directly visible: participants with higher CAD scores showed lower sorting consistency. Thus, even when the overall category structures remained close to that of trichromats, the fidelity of individual assignments was constrained by residual red-green sensitivity. Dichromats showed no comparable CAD relationship, most likely because the absence of one cone class imposes a more global constraint on the sensory evidence available for categorisations. Their responses therefore appear to rely more strongly on cues that remain available, especially lightness, together with learned regularities linking surface appearance to color terms. In this view, CVD does not erase categorical color representations; instead, it reduces the precision with which sensory inputs can be mapped onto otherwise stable categories.

### Category stability and sensory precision

The test-retest results provide a complementary view of the same mechanism. If color categories in CVD were merely unstable guesses, repeated sorting should have produced large changes across the two sessions. Instead, all groups showed substantial within-observer reliability, while stability declined systematically with the severity of the sensory deficit. As expected, trichromats showed the highest test-retest reliability, anomalous trichromats were somewhat less stable, and dichromats showed the largest reductions in stability. The same graded pattern was visible in the prototype selections: trichromats chose highly similar prototypes across the two sessions, anomalous trichromats showed only modest additional variability, whereas dichromats showed larger shifts while retaining a recognizable category structure.

This pattern suggests that color categories are still present in participants with CVDs, but that depending on the severity of the CVD the mapping from sensory input to category labels becomes less precise. In anomalous trichromats, this interpretation was supported directly by the values of the RG-CAD test: observers with higher RG-CAD thresholds, indicating poorer red-green discrimination, showed lower sorting consistency and reduced test-retest stability. Dichromats showed no comparable relationship between their RG-CAD scores and their sorting variability. All five dichromats had similarly high RG-CAD values, and the absence of one cone class imposes a global limit on discrimination and sorting. Their sorting nevertheless remained structured despite this greater variability. Divergences from the trichromat consensus were strongest for low-chroma chips, and the clearest category-level differences occurred for green and purple. Thus, reduced stability did not reflect random responding, but a less precise and more cue-dependent mapping of surface colors onto learned categories.

These results align with prior work showing that red-green CVD observers exhibit greater variability and uncertainty in color categories than normal trichromats [19,22,23,25,42,43]. Together, the present findings suggest that the categorical stability of color assignments depends on sensory precision: trichromats show highly stable color categories, anomalous trichromats show only modest reductions and remain largely comparable to trichromats, and dichromats show larger reductions in stability while still retaining core categorical structure.

### Enchroma

The EnChroma condition provided a complementary test of whether changing the spectral input by filters would immediately alter categorical color sorting. After approximately two hours of wearing the glasses, and while continuing to wear them during the subsequent sorting task, anomalous trichromats did not shift measurably toward the trichromatic patterns. Within-group agreement was essentially unchanged, and agreement with the trichromat consensus increased only slightly and non-significantly. Thus, in our test condition the use of EnChroma glasses did not change the ability to discriminate and sort color chips nor did it cause a large immediate reorganization of category assignments.

This result should be interpreted cautiously, because we tested only short-term exposure, with approximately two hours of adaptation, and cannot rule out effects of long-term adaptation. Nevertheless, it is consistent with the broader conclusion of the study: categorical color behavior is not determined solely by the momentary spectral input, but reflects relatively stable learned mappings between surface appearance and color terms. The result also aligns with recent objective evaluations reporting no or limited benefits of EnChroma glasses and related notch-filter aids for red-green CVD [44–53].

### Limitations

Several limitations of the present study should be acknowledged. First, although the stimulus set provided broad coverage of Munsell color space, it contained many saturated chips. This may have made some category assignments easier than they would be for less saturated or more ambiguous surface colors. However, the inclusion of additional low-chroma chips allowed us to test performance in less saturated regions, and our tested anomalous trichromats and dichromats still showed structured sorting and above-chance categorization for these stimuli. Thus, the main results are unlikely to depend solely on highly saturated chips. Here future work is needed to sample low-chroma and boundary regions even more densely.

Second, the restriction to 11 basic color categories may have constrained participants’ responses. This choice was mainly motivated by consistency with prior work, including large-scale studies supporting the prominence of these categories (e.g., [54]). It may have obscured finer hue distinctions that CVD participants might otherwise have drawn. Future work should therefore sample low-chroma and category-boundary regions more densely.

Finally, the relatively small number of dichromatic participants limits the extent to which subtype-specific conclusions can be drawn. While grouping protan and deutan observers is consistent with previous functional approaches, larger samples are required to examine systematic differences between these subtypes in greater detail.

## Conclusion

In summary, our sorting task shows that color categories in anomalous trichromats are highly stable and closely resemble those of trichromats across a broad set of physical Munsell samples. Dichromats showed reduced stability and greater divergence from trichromat categories, but nevertheless retained coherent and structured categorical organization. The principal effect of color vision deficiency is therefore not a collapse of color categories, but a graded reduction in the precision with which colors are assigned to different categories. These findings suggest that categorical color representations are not determined solely by sensory precision, but reflect learned and cognitive structures that remain surprisingly robust even when chromatic signals of the L- or M-cones are substantially degraded or lost.

## Acknowledgments

This project was supported by European Research Council ERC AdG Color 3.0 (884116), by the DFG Sonderforschungsbereich SFB TRR 135 (222641018) and by DFG Excellence Cluster EXC 3066/1 “The Adaptive Mind” (533717223). We thank all our participants for their patience during all experiments and color vision tests and their commitment to complete the whole data collection for this study.

## Data availability

All data and code will be made available on zenodo upon acceptance of the manuscript.

## Footnotes

1 Our results were first presented at the 2024 Vision Sciences Society meeting [28]. Hurlbert and colleagues have presented closely related work at conferences [29,30].

